# Nucleolar disruption, activation of P53 and premature senescence in POLR3A-mutated Wiedemann-Rautenstrauch Syndrome fibroblasts

**DOI:** 10.1101/2020.01.29.925131

**Authors:** Cindy Tatiana Báez-Becerra, Estefania Valencia-Rincón, Karen Velásquez-Méndez, Nelson J. Ramírez-Suárez, Claudia Guevara, Adrian Sandoval-Hernandez, Carlos E. Arboleda-Bustos, Leonora Olivos-Cisneros, Gabriela Gutiérrez-Ospina, Humberto Arboleda, Gonzalo Arboleda

## Abstract

Recently, mutations in the RNA polymerase III subunit 3A (POLR3A) have been described as the cause of the neonatal progeria or Wiedemann-Rautenstrauch syndrome (WRS). POLR3A have important roles in the regulation of transcription of small RNAs, including tRNA, 5S rRNA and U6 snRNA. We aim to describe cellular and molecular features of WRS fibroblasts. Cultures of primary fibroblasts from one WRS patient [monoallelic POLR3A variant c.3772_3773delCT (p.Leu1258Glyfs*12)] and one control were grown. Mutation in POLR3A causes a decreased in the expression of POLR3A mRNA and protein and a sharp increased of mutant protein. In addition, there was an increased in its nuclear localization. These changes were associated to an increase number and area of nucleoli, a significantly larger nuclear area, and a high increased in the expression of pP53 and pH2AX. All these changes were associated to premature senescence. The present observations add to our understanding of the differences between HGPS and WRS, and opens new alternatives to study cell senesce and human aging.

## 1. INTRODUCTION

Progeroid syndromes are a heterogeneous group of diseases characterized phenotypically by features of premature aging and *in-vitro* by premature cellular senescence (Martin, 1982; Sinha et al., 2014). The prototype progeria syndrome or Hutchinson-Gilford progeria (HGPS, OMIM 176670) is caused by a mutation in exon 11 (G608G) of the *LMN A/C* gene (De Sandre-Giovannoli et al., 2003; Eriksson et al., 2003). As a consequence, a cryptic splicing site generates an abonormal protein known as progerin, that produces typical cellular abnormalities most importantly at the nuclear level (Goldman et al., 2004; Mounkes and Stewart, 2004; Scaffidi and Misteli, 2005). Atypical progeroid syndromes have also been associated with mutations in *LMN A/C* (Csoka et al., 2004; Huang et al., 2005).

Wiedemann-Rautenstrauch Syndrome (WRS, MIM 264090) is a neonatal progeroid disorder (Arboleda et al., 2011; Paolacci et al., 2017), clinically characterized by signs of premature aging present at birth such as pseudohydrocephalus, craniofacial disproportion, triangular face, neonatal teeth, thin skin, reduction of subcutaneous fat and intrauterine growth retardation (Arboleda G. et al; Paolacci et al., 2017). WRS patients have a varied lifespan, ranging from months of age to adulthood (Arboleda and Arboleda, 2005; Paolacci et al., 2017). The neonatal phenotype is a key feature that differentiates WRS from HGPS. In HGPS the progeroid features appear in post-natal life (Ahmed et al., 2018; Piekarowicz et al., 2019; Ullrich and Gordon, 2015). In addition, no mutations have been found in the coding regions of *LMN A/C* in WRS (Cao and Hegele, 2003; Morales et al., 2009). Whether these two progeroid syndromes share a common aetiology based on cell abnormalities and/or molecular changes remains to be analyzed.

Recently, we and other groups demonstrated that WRS syndrome is caused by mutations in the RNA Polimerase III subunit A (*POLR3A*) gene (Jay et al., 2016; Paolacci et al., 2018). *POLR3A* mutations reduce the catalytic function of RNA polymerase III (POLR3) (Jay et al., 2016; Paolacci et al., 2018). Interestingly, allelic mutations in *POLR3A* gene has previously been reported to cause syndromes associated to hypomielinization, such as the hypomyelination, hypodontia and hypogonadotropic hypogonadism syndrome (4H syndrome) (Bernard et al., 2011; Saitsu et al., 2011). However, the exact role of POLR3A in cell senescence and aging is still unknown.

POLR3 is composed by 17 subunits with a molecular weight of 700kD, which is assembled in the cytoplasm and then translocated to the nucleus in a process mediated by chaperones (Han et al., 2018; Khatter et al., 2017). POLR3A is the largest subunit of the POLR3 complex. POLR3A is a 1390 amino acids (MW 156 kDa) protein that forms the catalytic core of the enzyme together with POLR3B (Han et al., 2018; Willis and Moir, 2018). Structurally, POLR3A is mainly form by alpha helixes conformations, that have been predicted to promote the movement of the polymerase along the interface RNA-DNA (Han et al., 2018). POLR3 is responsible for the transcription of genes coding for all transfer RNAs (tRNA), the 5S subunit of ribosomal RNA (5S rRNA), U6 small nuclear RNA (U6 snRNA), among others (Lesniewska and Boguta, 2017; Turowski and Tollervey, 2016; Willis and Moir, 2018). It also has been shown that POLR3A is important for the proper function of the nucleolus, ribosome assembly and protein translation that determines the metabolic state of the cell (Tiku and Antebi, 2018).

In the present study we aim to analyze several cellular and molecular changes in primary POLR3A-mutated WRS fibroblasts and compared them with human derived control and HGPS fibroblasts. We found that mutations in POLR3A cause an early senescent phenotype associated to alterations in the number and structure of nucleolus. The present observations add to our understanding of the cellular and molecular characteristics of WRS fibroblasts, and to the differences between HGPS and WRS. These observations opens new alternatives to study cell senescence and human aging nor currently known.

## 2. MATERIALS AND METHODS

### 2.1 Cell Culture

Primary human fibroblasts were obtained from forearm skin biopsies of a 13-year-old WRS patient [monoallelic *POLR3A* variant c.3772_3773delCT (p.Leu1258Glyfs*12)] (Paolacci et al., 2018). Control cells were obtained from a healthy female control of similar age to the patient. This study was approved by the Institutional Review Board of the Universidad Nacional de Colombia, according to the ethical standards laid down by the Declaration of Helsinki (1964) and its later amendments. All patients or legally authorized representatives of those unable to give consent for themselves gave their informed consent prior to their inclusion in the study. In addition, primary fibroblasts from one HGPS patient (AG01972, 1824 C-T, Coriell Institute) were also analyzed. Fibroblasts cultures were culture under standard conditions in MEM medium supplemented with 10% fetal bovine serum (Thermo Scientific, Rockford IL, USA, Ref. 12657) and 1% penicillin/streptomycin (Lonza, Basel, Switzerland, Ref. 17-603E) at 37°C in a humidified 5% CO2 incubator. Cellular analysis was performed at population passages described in each figure.

### 2.2 In-silico protein structure analysis of the POLR3A mutation

Alterations in the structure of the human POLR3A associated to the c.3772_3773delCT (p.Leu1258Glyfs*12) mutation were analyzed by homology modeling based in the protein data bank (PDB) structure 5fj8 of the Saccharomyces cerevisiae Pol III elongating complex at 3.9 Å resolution and the Pol III enzyme in two different conformations at 4.6 and 4.7 Å (Hoffmann et al., 2015). The structure analysis of the POLR3A mutation was done using the CCP4MG software (McNicholas et al., 2011), located in the chr10:79741306, which generates a stop codon located 12 amino acids from the mutation site. Amino acid sequence alignment between diverse species was done by using the Geneious prime software (V. 8.1; https://www.geneious.com).

### 2.3 RT-qPCR

WRS patients and healthy control fibroblasts were growth in 100 mm Petri dishes. RNA was extracted with TRIzol™ Reagent (ThermoFisher Scientific, Rockford IL, USA) according to manufacturer’s protocol. 100 μL of TRIzol™ Reagent was used to 10 mm Petri dish. An equal volume of chloroform was added and then, samples were shaken by vortex for 2 minutes until get an emulsion. Aqueous phase was recovered and then RNA was precipitated with isopropanol. RNA was rinsed with 80% ethanol twice and then quantify in NanoDrop^TM^ Spectrophotometer (ThermoFisher Scientific, USA). 200 to 3000 ng/μL RNA was recovered in WRS samples and control. Fresh extracted RNA from MO3.13 cells was used as positive control to curve standard. 200 ng/μL was used for subsequent reverse transcription assay with Luna^®^ Universal One Step RT-qPCR kit (E3005, New England Biolabs) according manufacturer’s protocol. Each reaction was made in a 10 μL final volume. RT-qPCR was made in a CFX96 Touch^TM^ Real-Time PCR Detection System (BioRad) with 48 cycles. Primers used were: POLR3A-WT forward: 5’-AAGCTTCTGGTGGAAGGTGA-3’, reverse: 5’-CTCCTGTCGATGCTCATGC-3’; POLR3A-MUT forward: 5’-GAAAGAGGATCTCCCCAAGG-3’, reverse: 5’-CGCAGGTTATCACCTTCCAC-3’; 5SrRNA forward: 5’-GCCATACCACCCTGAACG-3’, reverse: 5’-AGCCTACAGCACCCGGTATT-3’; tRNA-Leu-CAA forward: *5’-*CTCAAGCTTGGCTTCCTCGT-3’, reverse: *5’-* GAACCCACGCCTCCATTG-3’; 7SK RNA [13] forward: 5’-AGAGGACGACCATCCCCGAT-3’, reverse: 5’-TGGAAGCTTGACTACCCTACGT-3’; 18S rRNA forward: 5’-GTAACCCGTTGAACCCCATT-3’, reverse: 5’-CCATCCAATCGGTAGTAGCG-3’; 28S rRNA, forward: 5′-AGAGGTAAACGGGTGGGGTC-3′, reverse: 5’-GGGGTCGGGAGGAACGG-3’ (Macrogen, South Korea) and beta-actin forward: 5’-CCTGGTCGGTTTGATGTT-3’ and reverse: 5’-GTGCGACGAAGACGA-3’ (Invitrogen, USA). Data analysis was made in CFX Manager^TM^ Software (BioRad) and GraphPad^TM^ Prism 5.0 (La Jolla, California, USA). ΔΔCt method were used to relative quantification. Other methods as Standard curve and Livak were used to confirm results.

### 2.4 Western Blot

Protein expression levels were evaluated by western blot. Cell cultures at 70% of confluence were scraped, pelleted and lysed at 4 °C for 10 min and ultrasound at 20% amplitude in lysis buffer (Ripa buffer, Thermo scientific) containing 1% protease inhibitor cocktail (Complete Mini-Roche Molecular Biochemicals; Mannheim, Germany) and 1% phosphatase inhibitor cocktail (Phospho-STOP Roche Molecular Biochemicals; Mannheim, Germany). Lysates were centrifugated at 13000 rpm at 0°C, then protein was quantified using BCA protein assay kit (Thermo Scientific, Rockford IL, USA, Ref. 23227) using bovine serum albumin (BSA, Merck, Darmstadt, Germany, Ref.A2153) as the standard. SDS-PAGE was made with 4–20% Mini-PROTEAN^®^ TGX™ Precast Protein Gels 10-well (BioRad, 4561094) with 30 μg of protein lysate for well and running was 70 V for 2 hours. Protein transfer from gel to membrane was performed with a 0.45 μm PDVF Amersham^TM^ Hybond^TM^ membrane (GE Healthcare, formerly Amersham Biosciences, and Piscataway NJ, USA, Ref.40600023) previously activated with absolute methanol. The transfer was made at 4 °C overnight. Membrane was blocked with blocking buffer consisting of 5% BSA or 5% fat-free milk in 1X Tris Saline Buffer (TTBS) pH 7.6. Then PDVF membranes were incubated in primary antibodies [anti-Lamin A (1:1000, Cell Signalling, Danvers, MA, USA, Ref. 2032), anti-POLR3 (1:500, Novus Littleton CO, Ref. NBP1-83204), anti-P16 antibody (1:1000, Cell Signalling, Danvers, MA, USA, Ref. 80772), anti-phospho P53 (Serine 15; 1:1000; Cell Signalling, Danvers, MA, Ref. 9284L), anti-phospho H2AX (Serine 139; 1:1000, Cell signaling, Danvers, MA, Ref. 2577S)] in blocking buffer at 4°C overnight. β-actin anti-mouse antibody (1501L, Millipore) was used at 1:2000 dilution in blocking buffer. Then the membrane was washed 3 times with TTBS. Secondary antibodies were used at 1:2000 dilution in blocking buffer: Anti-rabbit IgG HRP-linked (7074S, Cell Signaling) and anti-mouse IgG HRP-linked (7076, Cell Signaling), for one hour at room temperature. Peroxidase activity was evaluated by Novex™ ECL Chemiluminescent Substrate Reagent Kit (ThermoFisher Scientific, Rockford, IL, USA, Ref. WP20005) and the image acquired using ChemiDoc^TM^ Imaging Systems with Image Lab^TM^ Software (BioRad, Hercules, CA, USA). The images were analyzed with Image Lab software (Bio-Rad, Hercules, CA, USA) and Prism5-GraphPad (San Diego, CA, USA). Densitometric analysis was normalized with respect to beta actin.

### 2.5 Immunofluorescence

Primary fibroblast was seeded on coverslips covered with poli-L Lysine (Merck, Darmstadt, Germany, Ref. P1274) and grown at 70% of confluence. The cells were fixed with 4% paraformaldehyde containing 4% sucrose in PBS 1X at room temperature for 15 minutes. Cells are then permeabilized with 0.1% Triton X-100 solution in Tris Buffered Saline (TTBS) for 15 minutes, and blocked with blocking buffer (0.5% FBS, 0.1% BSA in TTBS) for 30 minutes, followed by incubation with primary antibodies [anti-POLR3a (1:500; Novus Littleton CO, Ref NBP1-83204), anti-fibrillarin (1:250; Cell Signalling, Danvers, MA, Ref. 2639S), anti-phospho P53 Serine 15 (1:100; Cell Signalling, Danvers, MA, Ref. 9284L), anti-phospho H2A (1:100; Cell Signalling, Danvers, MA, Ref. 2577S)]. Primary antibodies were incubated overnight at 4°C in blocking buffer (5% SFB and 1% BSA in TTBS). Next, coverslips were incubated for 2 hours in secondary antibody [Goat-anti-rabbit (1:3000; Alexafluor 568, Invitrogen, ThermoFisher Scientific, Rockford IL, USA, Ref. A11011)]. Alexa Fluor® 488 Phalloidin (Invitrogen, ThermoFisher Scientific, Rockford IL, USA) was used to stain cytoskeleton and 1 μM Dapi (sc-3598, ChemCruz^TM^ Biochemicals, USA) was used as a counterstain. Finally, coverslips were mounted onto a slide with fresh mounting media (70% glycerol in PBS 2X) and sealed with colorless nail polish and viewed with a Nikon Eclipse C1 Plus Ti (Tokyo, Japan) scanning confocal fluorescence microscope with 60X magnitude. Samples were stored at 4°C for further analysis. All image quantification was performed with ImageJ software (National Institutes of Health, USA). DNA was counterstained with DAPI. Two-tailed student tests were used to calculate P-values.

### 2.6 Beta-Galactosidase analysis

Activity of lisosomal Beta-galactosidase (B-gal) can be detected *in situ* in most mammalian cells by citochemistry assays using a chromogenic substrate 5-bromo-4-cloro-3-indolyl galactopyranoside (X-gal) (Kuo and Wells, 1978). It has been demostarted that B-gal activity at pH 6 is specifically associated to cell senescence (Childs et al., 2019; Dimri et al., 1995). B-gal activity was measure in fibroblasts from WRS, HGPS and control cells. Briefly, cells were seeded in 24 well-plates, washed 3 times in DPBS (Dulbeccos Phosphate Buffered Saline), fixed in 2% (v/v) formaldehyde (Sigma, USA) washed again in DPBS and stained with 1mg/ml X-gal (5-bromo-4-chloro-3-indolyl-β-galactopyranoside, Promega, Madison, WI, US) in DMF (Dimetil formamide, Mallinckrodt, St Louis, MO, USA) plus 0,1M 4 ml citric acid and sodium phosphate (0,2M) at pH 6,4. Two controls were used: pH 7,4 (as negative B-gal) and pH 4 (as positive B-gal). Cells were then grown at 37°C without CO2 for 24 hours. Analysis of cells was done for 5 fields and the number of blue cells was quantified under an inverted NIKON TE600 microscope (Childs et al., 2019; Dimri et al., 1995).

### 2.7 Telomere length analysis

Genomic DNA was extracted from primary fibroblasts using a commercial purification kit (Promega, Madison, WI, USA). DNA was store in TE (10 mM Tris, pH 7.5, 1 mM EDTA) at 4°C until use. Telomeric repets were calculated as described (Gil and Coetzer, 2004a, b; Vasilishina et al., 2019) using a LightCycler 2,0 (Roche Diagnostics, Basilea, SWI). Briefly, 2ul of DNA (20 ng of DNA) was added at 10ul of a reaction buffer (Light Cycler Fast Start DNA Master SYBR Green Kit, Roche Diagnostics, Basilea, SWI) and 1% SDS. Relative telomere length was normalized with amplification of single-copy gen 36B4, which codifies for a ribosomal phosphoprotein PO (chromosome 12). Primers used were described previously (Cawthon, 2002): Telomeric primers: forward: 5’CGGTTTGTTTGGGTTTGGGTTTGGG TTTGGGTTTGGGTT-3′ and reverse: 5’GGCTTGCCTTACCCTTACCCTTACCCTTACCCTTACCCT-3′; One copy gen: (36B4): 36B4u, CAGCAAGTGGGAAGGTGTAATCC; 36B4d, CCCATTCTATCATCAACGGGTACAA. SH-SY5Y and HT1080 telomerase positive cell lines were used as a positive control (Binz et al., 2005; Villa et al., 2004). RT PCR was carried out as follows: initial denaturation at 95°C for 10minutes; 30 cycles of denaturation at 95°C for 5 seconds, annealing at 56°C for 10 seconds, extension at 72°C for 60 seconds, for telomeric repeats. For one copy gen 40 cycles at 95°C for 5 seconds, annealing at 58°C for 10 seconds and extension at 72°C for 40 seconds (Gil and Coetzer, 2004b).

### 2.8 Statistics

All values are presented as the mean ± SEM. Statistical significance between groups is determined using a student’s t for normal distributions and Mann-Whitney U for non-normal distributions (PRISM 5 GraphPad, San Diego, CA, USA).

## 3. RESULTS

### 3.1 Structural analysis of PolR3A mutation in the present case

Protein structure analysis of yeast POLR3 (RPC1) shows that POLR3A is at the center of interactions of different sub-units of POLR3 that are important for proper POLR3 function (Figure 1A). The c.3772_3773delCT (p.Leu1258Glyfs*12) mutation associated to the present WRS case creates a premature stop codon (Figure 2C: *) and lost of 121 amino acids at the C-terminal domain of the protein that is predicted to generate a POLR3A truncated protein of ∼142 KDa (Figure 1B and C). The deleted C-terminal region at the RNA_pol_Rbp1_5 domain of POLR3A (in red: Figure 1A and B) is important for its proper interaction with POLR3B, POLR3D, POLR3E, POLR3F and POLR3G isoforms (Figure 1A). Amino acid sequence analysis of POLR3A surrounding the c.3772_3773delCT (p.Leu1258Glyfs*12) mutation shows a high degree of conservation in different species (Figure 1C).

**Figure 1.**
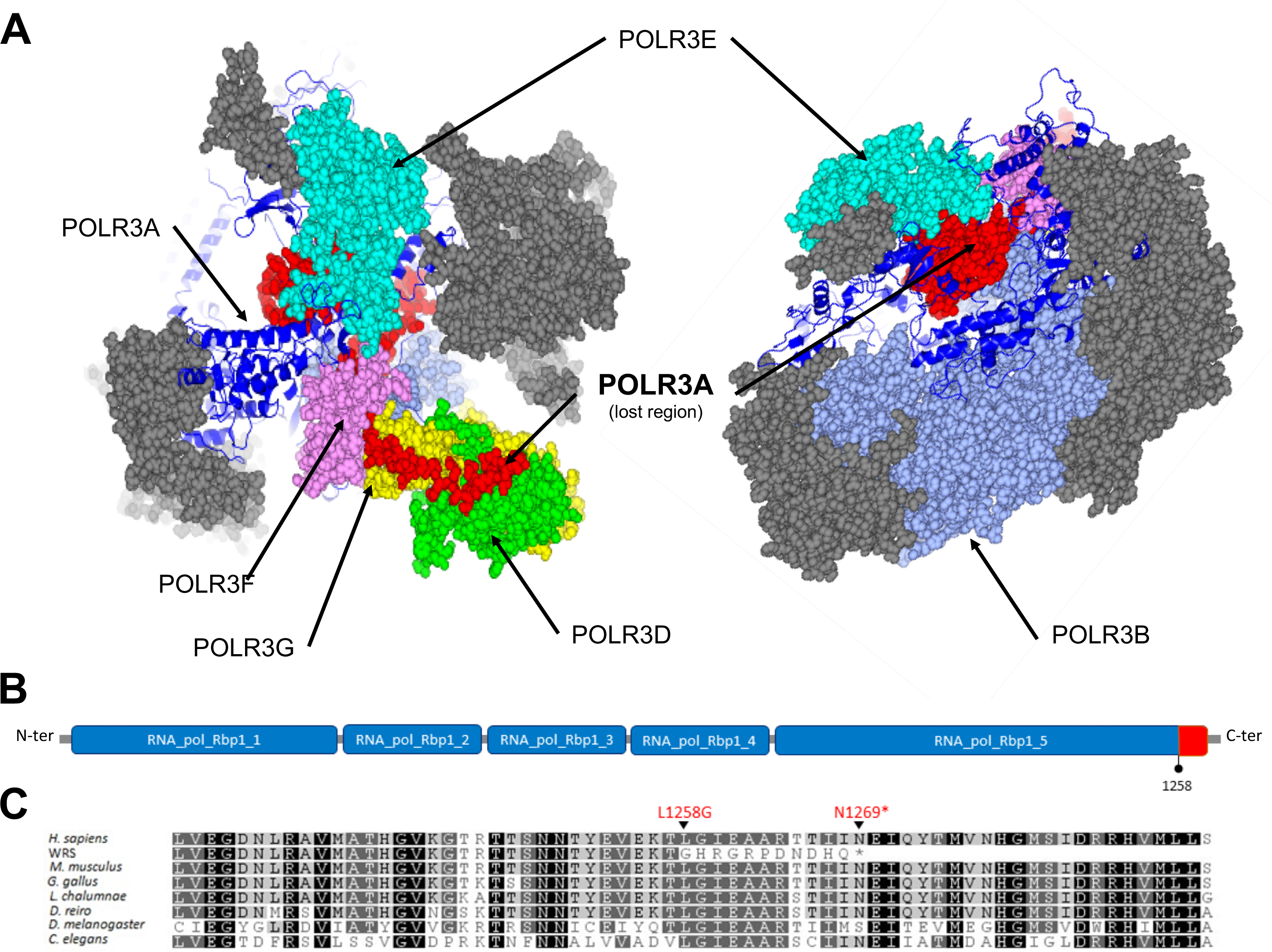
Protein structure analysis of the C-terminal region of POLR3A c.3772_3773delCT (p.Leu1258Glyfs*12) mutation. (A) Protein structure of yeast POLR3 (RPC1) showing the interaction of different sub-units of POLR3 (POLR3A, POLR3B, POLR3D, POLR3E, POLR3F, POLR3G) (Hoffmann et al., 2015). POLR3A is shown in blue ribbons and the lost region of POLR3A associated to the c.3772_3773delCT (p.Leu1258Glyfs*12) mutation is shown in red spheres. Structure analysis was done using the CCP4MG software. (B) Domain structure of POLR3A showing the region deleted by the c.3772_3773delCT (p.Leu1258Glyfs*12) mutation at the C-terminal region (in red), which is important for proper interaction of POLR3A with POLR3B, POLR3D, POLR3E, POLR3F and POLR3G isoforms. (C) Amino acid alignment of POLR3A homologous of different species showing a high degree of conservation. Shown is the surrounding amino acid sequence of the present WRS case associated to the c.3772_3773delCT (p.Leu1258Gly) mutation. Deletion in the present case causes the creation of a premature stop codon (N1269*) located 12 amino acids from the mutation site and lost of 121 amino acids at the C-terminal domain of the protein. The deletion generates a truncated POLR3A protein of 1269 amino acids (MW ∼142 KDa).

**Figure 2.**
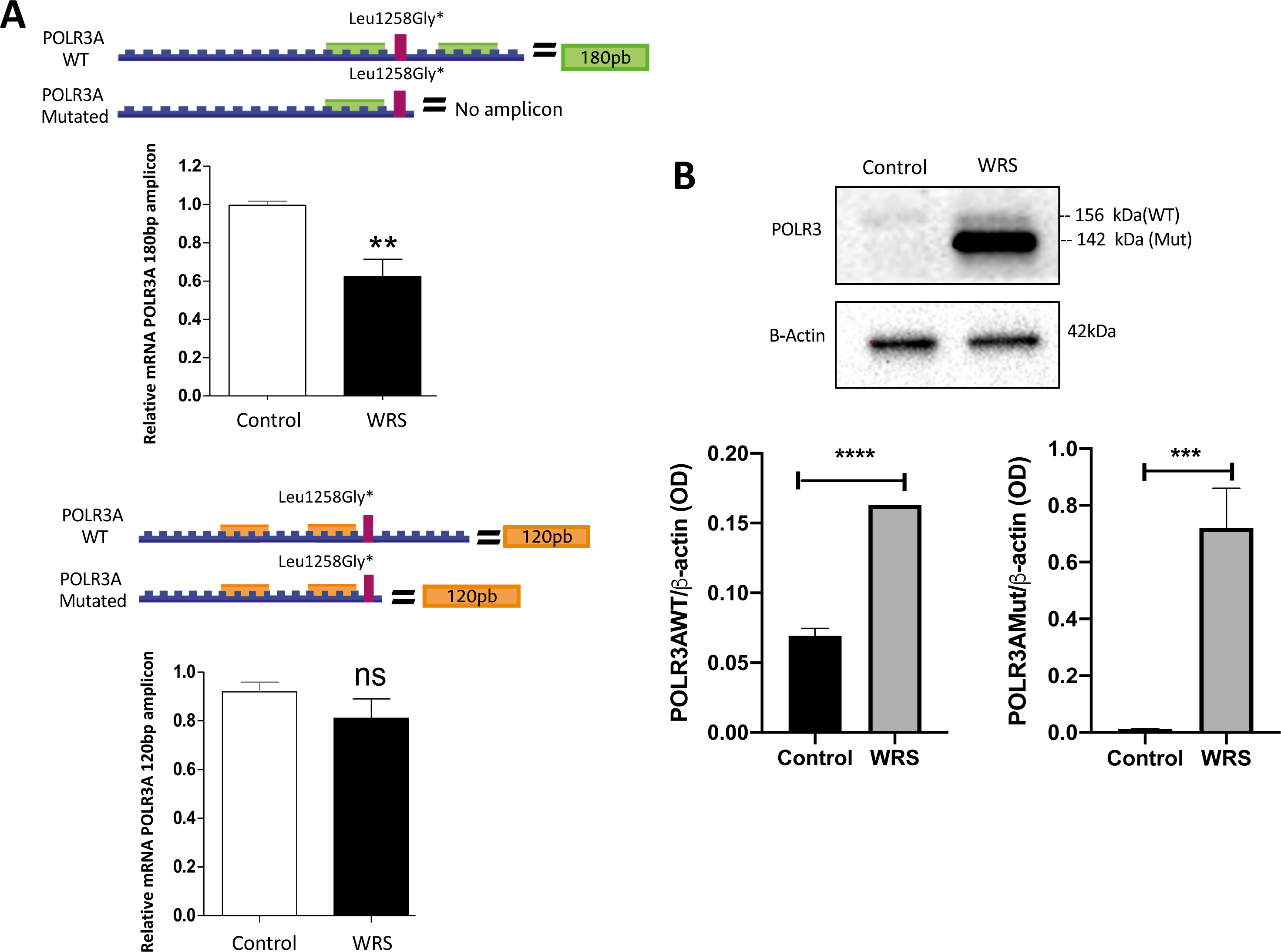
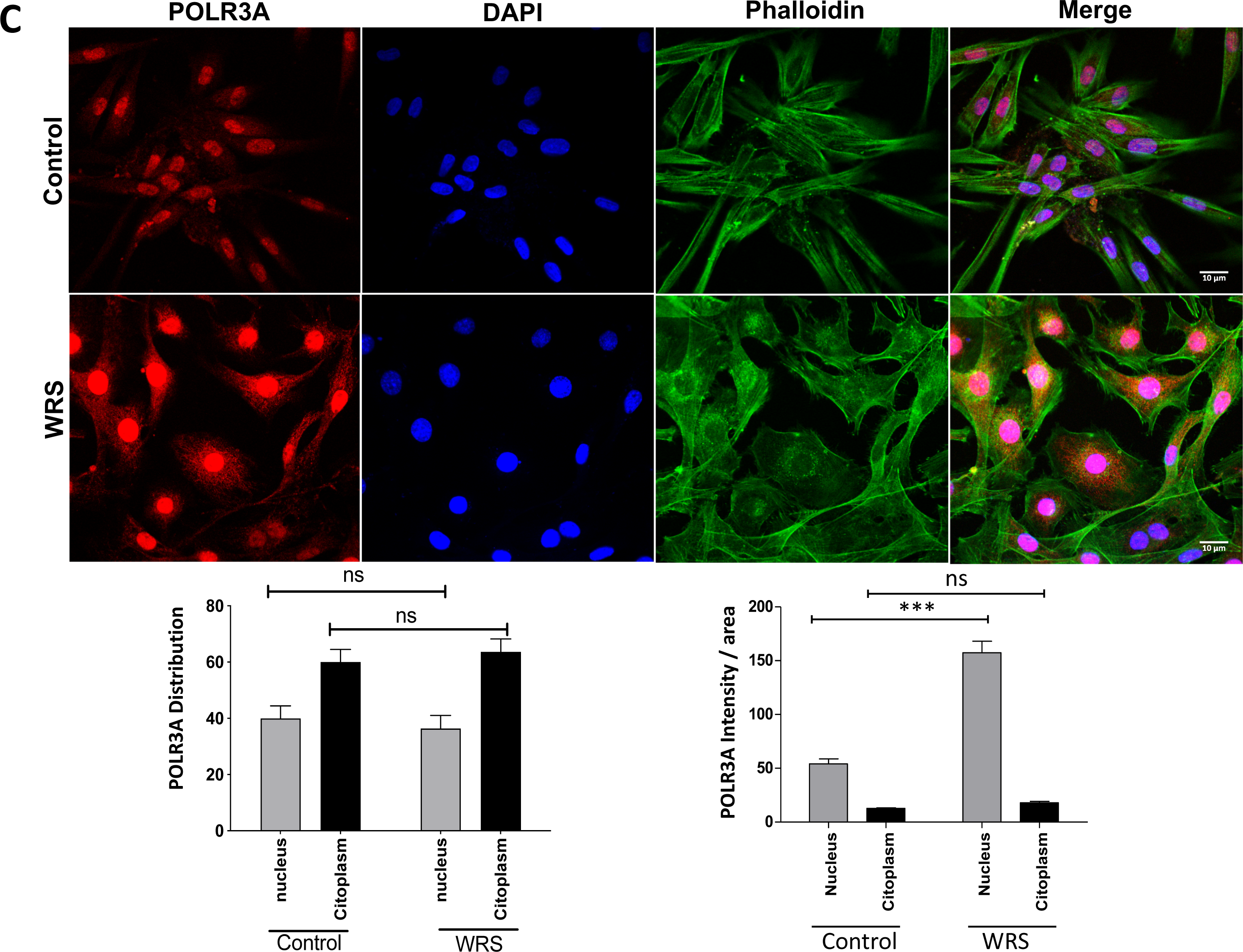
POLR3A expression and compartmentalization in WRS fibroblasts. Expression of POLR3A was analyzed by (A) RT-PCR, (B) Western Blot and (C) immunohistochemistry, in primary human fibroblast from control and WRS. (A) RT-PCR analysis using two different sets of primers: one set of primers includes the mutation site and generates a 180bp amplicon; and a second set of primers that excludes the mutation site and generates a 120bp amplicon. There is a decreased in the content of the 180bp amplicon in WRS cells as compared to control cells; however, there are no differences in the expression of the 120bp amplicon between control and WRS cells. (B) In control cells there is a unique POLR3A band (156 KDa), while in WRS cells there are two bands of POLR3A protein corresponding to the normal (156 KDa) and the mutated isoform (∼142 KDa), both of which are highly increase in WRS fibroblsts as compared to control cells. The results are a mean of three independent experiments with n=3 in each case (***: p<0.001). (C) Immunostaining of POLR3A in control cells and WRS cells. In control and WRS cells localization of POLR3A is observed in the nucleus and cytoplasm, with a similar pattern of distribution. However, analysis of POLR3A expression related to nuclear and cytoplasmic area showed that POLR3A intensity increases in the nucleus of WRS as compared to control cells (ns: not significant; ***: p<0.001).

### 3.2 Analysis of POLR3A expression in WRS fibroblasts

POLR3A expression in WRS fibroblasts was analyzed by RT-PCR (Figure 2A) and western blot (Figure 2B). There is a reduction in the level of the 180pb POLR3A mRNA amplicon (Figure 2A, upper panel), although there are no changes in the expression of the 120kb POLR3A mRNA amplicon (Figure 2A, lower panel). These suggest a decreased in the normal mRNA and no changes in the total expression of POLR3A mRNA. POLR3A protein expression in control cells exhibit a band that correspond to the normal POLR3A isoform (165 KDa); in WRS cells there are two bands that correspond to the normal POLR3A (165 KDa) and a sharp increase (ten times fold increase) of a shorter band that corresponds to the mutated isoform (142 KDa) of the POLR3A protein (Figure 2B).

In addition, POLR3A immunohistochemistry studies showed that control and WRS cells have a similar pattern of distribution between nucleus and cytoplasm; however, in WRS cells there is a sharp increased in the intensity of the nuclear staining per area analyzed (Figure 2C). These changes suggest that the increased in the expression of the mutated POLR3A in WRS cells is trnalated into an accumulation of the mutant protein into the nucleus. This can compromise the function of the remaining normal protein most probably by altering its normal compartmentalization and by a dominant negative effect (Herskowitz, 1987).

### 3.3 The genes transcribed by RNA polymerase III (5S RNA, tRNA and 7SK) and ribosomal transcripts are deregulated in WRS fibroblasts

To analyze the effect of the c.3772_3773delCT and c.3G> T mutations in the transcriptional control exerted by POLR3A, the reporter genes 5SrRNA, tRNA-Leu-CAA isoaceptor and 7SK RNA were analyzed by RT-qPCR (Figure 3A). The effect of mutations in the POLR3A gene is also seen in the relative amount of 18S and 28S ribosomal transcripts. Quantification of the number of transcripts of the 18S and 28S eukaryotic ribosomal subunits was performed based on the data obtained by the RT-qPCR analysis (Figure 3B). Low levels of expression of the 18S and 28S subunits, which are part of the minor subunit of the ribosomal complex, are observed in WRS. In general, it is observed that the mutations analyzed in the *POLR3A* gene cause a decrease in the mRNA expression of POLR3A target genes. The alterations in expression of POLR3A-regulated genes may have important consequences in ribosomal biogenesis, protein synthesis and regulation of other polymerases such as RNA polymerase II (Canella et al., 2010).

**Figure 3.**
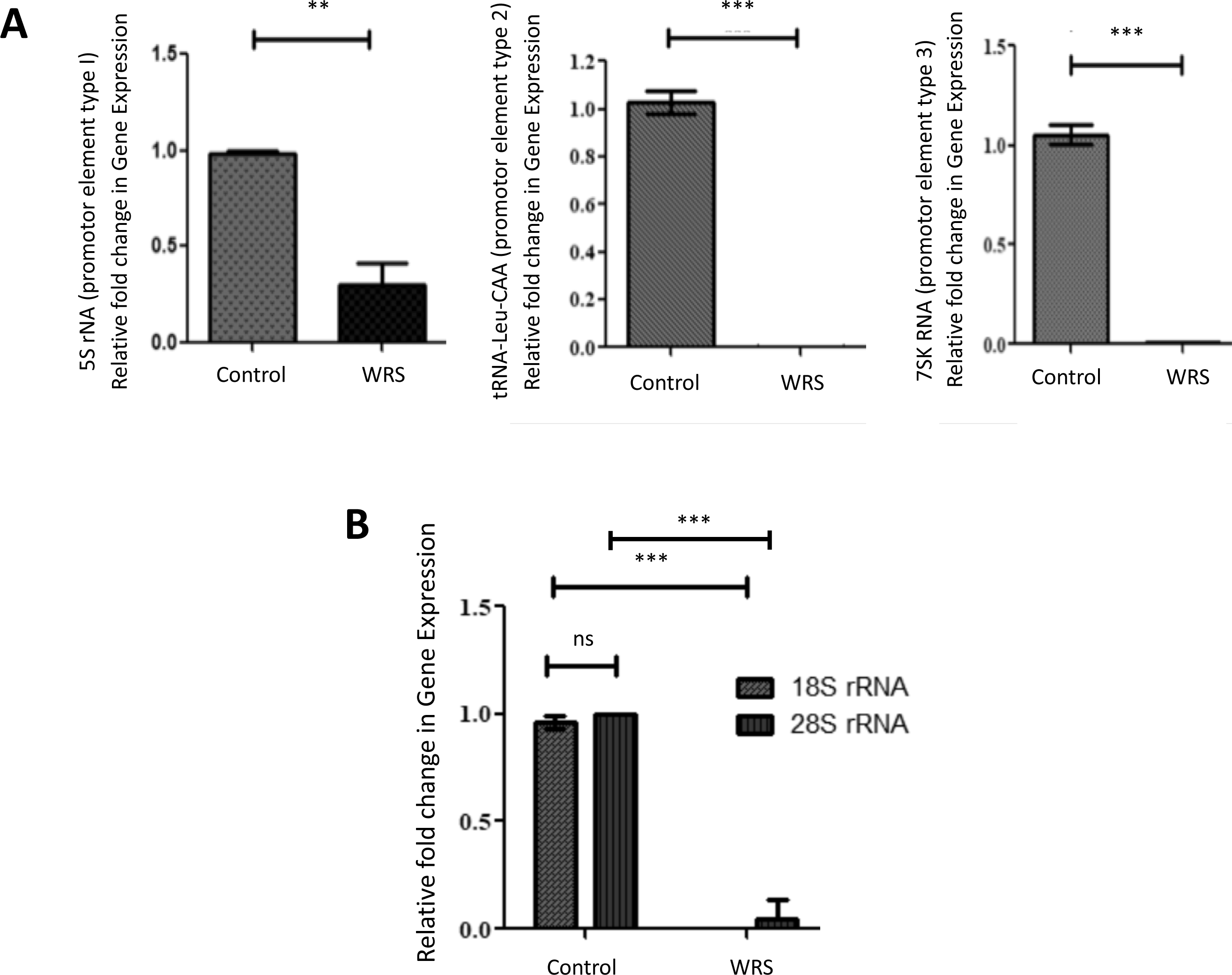
Expression of transcripts regulated by POLR3. (A) RT-qPCR quantification of 5S RNA, tRNA Leu CAA and 7SK RNA transcripts in control and WRS fibroblasts. There is a highly significant decrease in the expression level of mRNA of the genes analyzed in WRS cells as compared to control cells. Data were normalized to β-actin housekeeping gene (*: p<0.01; **: p<0.05; ***: p<0.001; ns: not significant). (B) Comparison of endogenous levels of the 18S and 28S ribosomal subunits between control and WRS fibroblasts. WRS cells showed a clearly significant decrease in the expression of 18S and 28S mRNA as compared to control cells (*: p<0.01; ***: p<0.001; ns: not significant).

### 3.4 Nucleolus are increase and disrupted in a WRS patient with the POLR3A c.3772_3773delCT mutation

The number and size of nucleoli varies from one cell type to another depending on transcriptional activity (Nemeth and Grummt, 2018). The main function of nucleoli is associated to control of ribosome biogenesis, which takes place in two different sub-structures: 1) the granular component, where pre-ribosomal assembly takes place; 2) the fibrillar center, rRNA transcription and dense fibrillar component, where pre-rRNA processing occurs. The nucleolus is usually labeled with nucleolin and its structures with NMP1, UBF and Fibrillarin. It has been observed that labeling of nucleolin and Fibrillarin are directly correlated (Baran et al., 1995).

By using Fibrillarin staining, we analyzed morphology and number of nucleoli (Figure 4A-D). WRS cells have an increase in the number (Figure 4B) and staining intensity (Figure 4D) of nucleoli per nucleus as compared to control fibroblasts. However, the area per nucleolus is significantly reduced in WRS cells (Figure 4C). In addition, the normal Fibrillarin staining observed in control cells (Figure 4A, bottom left panel) is clearly altered in WRS cells suggesting that there is disruption of nucleolar morphology (Figure 4A, bottom right panel).

**Figure 4.**
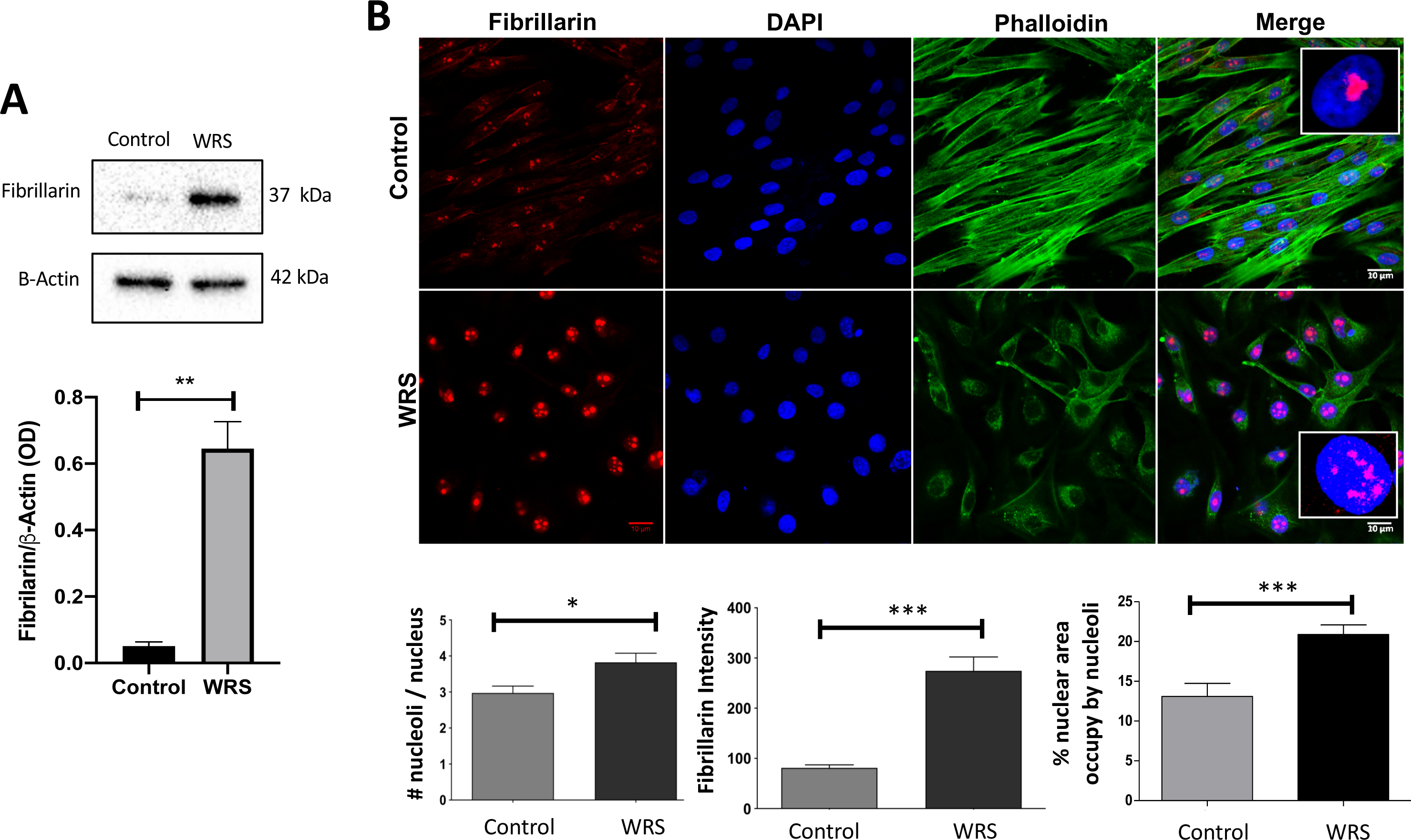
WRS fibroblasts showed disruption of nucleolar morphology. (A) Western blot analysis of Fibrillarin protein expression, showing a highly significant increase in its expression in WRS cells as compared to control cells. (B) Immunostaining of fibrillarin (nucleolus: red), DAPI (nucleus: blue) and phalloidin (cytoplasm: green) in control and WRS cells. There is a significant increase in the number and intensity of fibrillarin in WRS as compared to control cells. In addition, the percent of nuclear area occupied by Fibrillarin is also higher in WRS cells. In control cells fibrillarin staining is confined in round structures that correspond to normal nuclei morphology, while in WRS cells Fibrillarin staining is disrupted in multiple foci (as shown in the magnification insert). The results are a mean of 30 nuclei for each cell group (*: p <0,01; ***: p <0,001).

### 3.5 Increase in P53 and H2A expression in WRS fibroblasts

DNA damage leads to activation of a DNA damage response (DDR) that involves the activation of H2AX and P53 (Ou and Schumacher, 2018; Rodier et al., 2007). The H2AX histone is rapidly phosphorylated at serine-139 (gammaH2AX) in response to a broad range of DNA lesions and triggers subsequent events that have a central role in repairing DNA damage (Kopp et al., 2019). WRS fibroblasts showed strong phosphorylation of H2X (Ser139) as compared to control cells (Figure 5A), associated to an increased nuclear staining of H2AX (Ser139) (Figure 5B). Activation of P53 requires fine-tune posttranslational modifications, including phosphorylation at different sites (MacLaine and Hupp, 2011). Phosphorylation of P53 at position serine 15 (pP53-Ser15) leads to a reduced interaction between p53 and its negative regulator, MDM2, favoring the accumulation, stabilization and function of the P53 protein (Shieh et al., 1997). WRS cells display a highly significant increased in the level of pP53 (Ser15) (Figure 5C), associated to higher nuclear staining of pP53-Ser15, as compared to control cells (Figure 5D).

**Figure 5.**
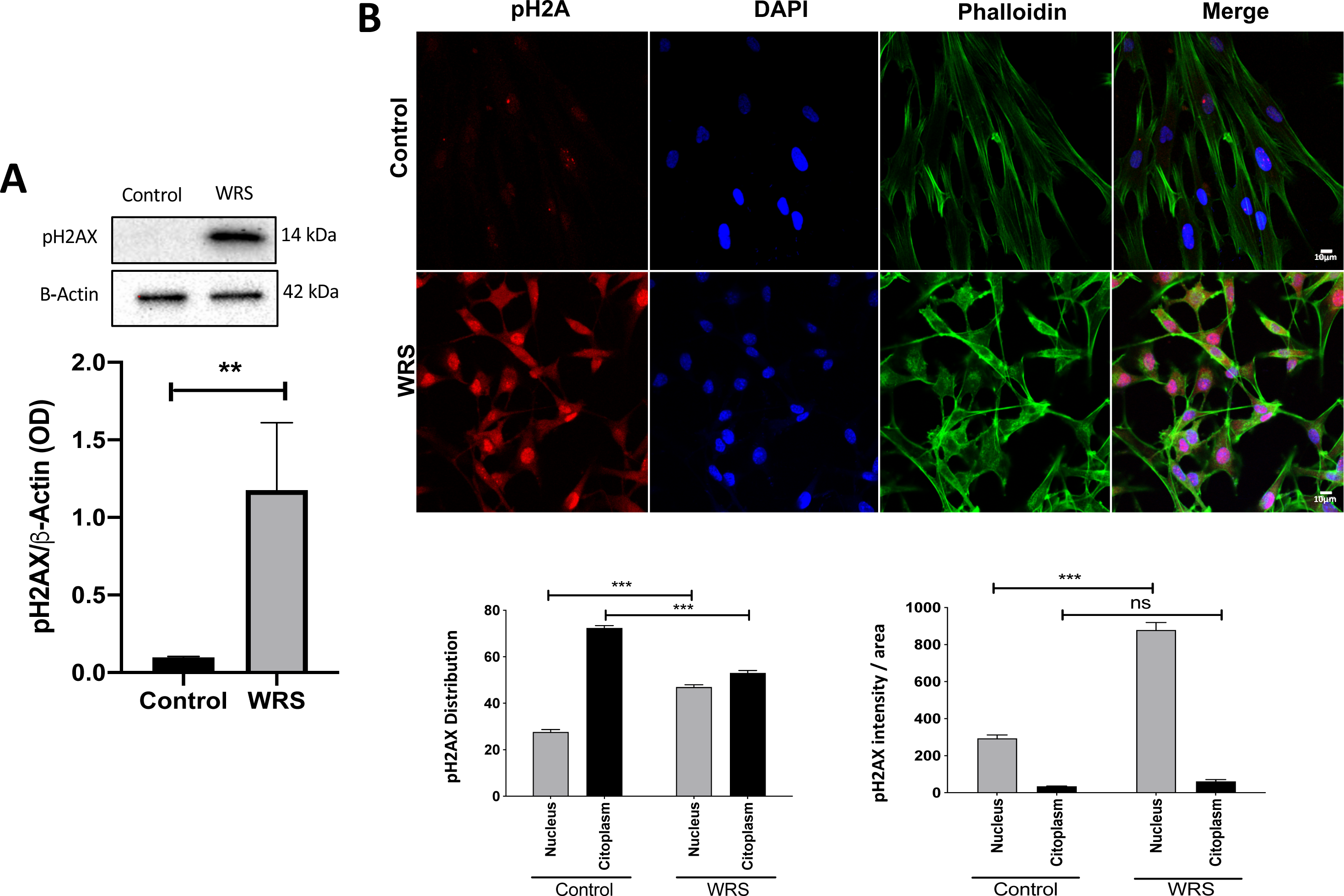
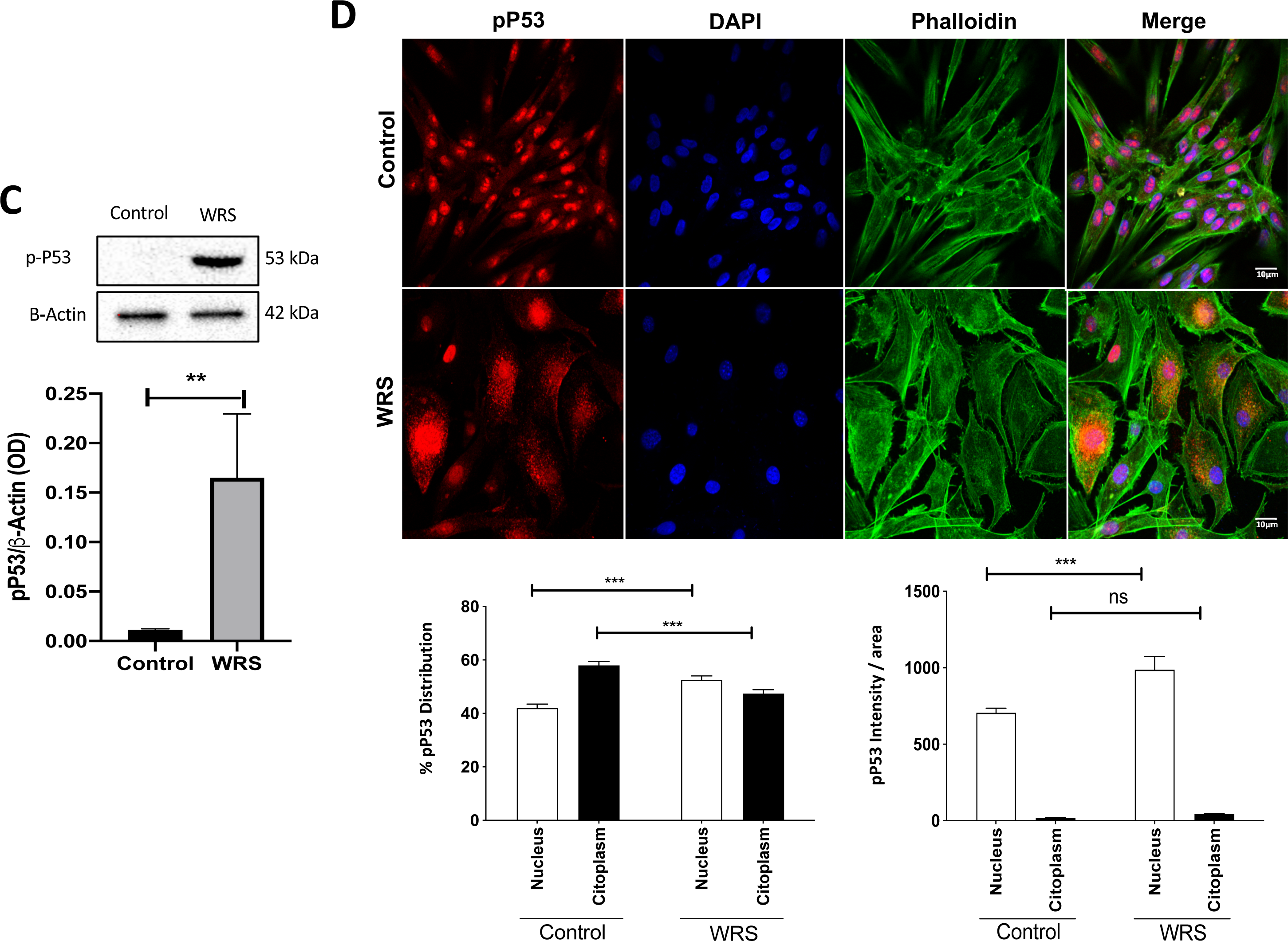
Increased expression of phospho H2AX (pH2AX) and phospho P53 (pP53) in WRS fibroblasts. (A) WRS fibroblasts showed an increase in the expression of phosphorylated H2AX (pH2AX) as compared to control fibroblasts; (B) Immunostaining of pH2AX (red), DAPI (nucleus: blue) and phalloidin (cytoplasm: green) in control and WRS cells. There is a significant increase in the nuclear staining of pH2AX and reduction in cytoplasmic staining in WRS as compared to control cells. In addition, there is also an increase in the intensity per area of pH2AX staining in the nucleus in WRS cells. (C) Expression of P53 phosphorylated at position serine 15 (pP53-Ser15) as analyzed by western blot. WRS fibroblasts showed an increase in the expression of pP53 as compared to control fibroblasts; (B) Immunostianing of pP53 (red), DAPI (nucleus: blue) and phalloidin (cytoplasm: green) in control and WRS cells. There is a significant increase in the nuclear staining of pP53 and reduction in cytoplasmic staining in WRS as compared to control cells. In addition, there is also an increase in the intensity per area of pP53 staining in the nucleus in WRS cells. The results are a mean of n=30 nuclei for each cell group (***: p <0,001; ns: not significant).

These observations suggest that in WRS fibroblasts have an increase in genomic damage that switches-on a H2AX- and P53-mediated DNA damage response (DDR) (Green and Kroemer, 2009; Kruiswijk et al., 2015).

### 3.6 Changes in nuclear morphology in WRS cells

Alterations in nuclear morphology are typical of different types of progeria (Broers et al., 2006; Mounkes and Stewart, 2004). Analysis of morphological characteristics of the nucleus showed that WRS nuclei are significantly larger in area and perimeter (Figure 6A, B and C). Additionally, it is observed that WRS nuclei have no changes in circularity (Figure 6D), have increase roundness (Figure 6E) and decreased aspect ratio (Figure 6F). These analyses suggested that WSR nuclei are in general larger and rounder.

**Figure 6.**
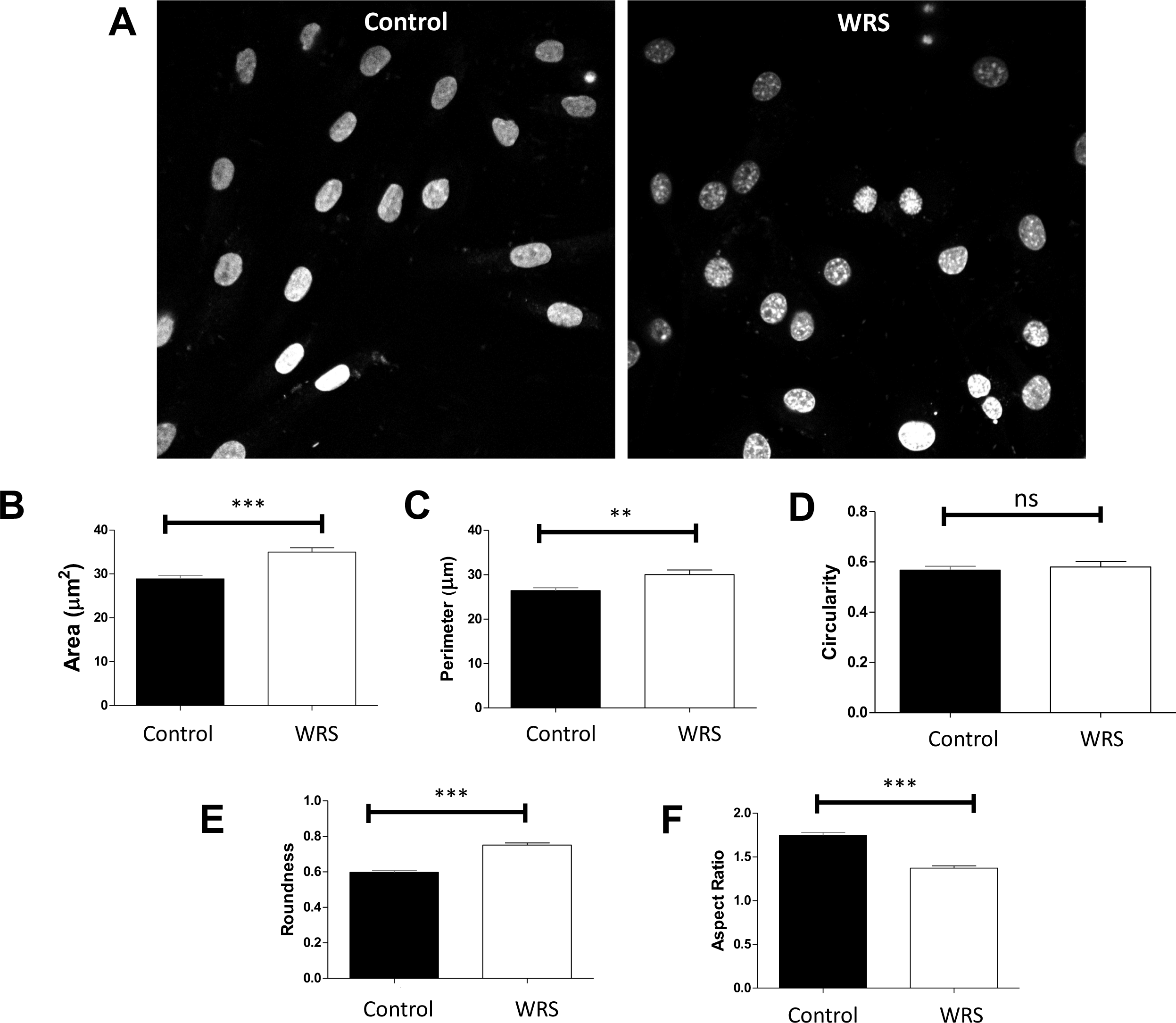
Increased size of nucleus in WRS fibroblasts. Analysis of nuclear morphology was performed by using confocal microcopy of control cells and WRS fibroblasts. (A-C) WRS nuclei are bigger than control nuclei as shown by significant increased in area and perimeter; and are rounder as shown by increase (E) roundness and (F) decrease aspect ratio of nucleus (relationship between greater diameter and smaller diameter, where values greater than 1 indicate elliptical shape and values equal to 1 indicate a perfect circle). These parameters were evaluated in 150 nuclei per culture by using ImageJ software. The comparisons were made with the Mann-Whitney test with (ns: not significant; ** p <0,01; *** p <0,001).

### 3.7 Comparative analysis of senescence markers, expression of Lamin A/C isoforms and telomere length between HGPS, WRS, and control cells

WRS cells underwent a process of cell senescence as determined by the expression of beta-galactosidase and P16, and decrease telomere length (Figure 7A, B and G). Beta-galactosidase staining was evident at population doubling (PD) 36 in WRS, while at PD 27 in HGPS and in controls at PD45 (Figure 7A). These changes were directly correlated with expression of P16, which was initially evident at PD 36 in WRS. Control cells did not showed expression of P16 even at PD 40, while HGPS showed P16 expression as early as PD25 (Figure 7B).

**Figure 7.**
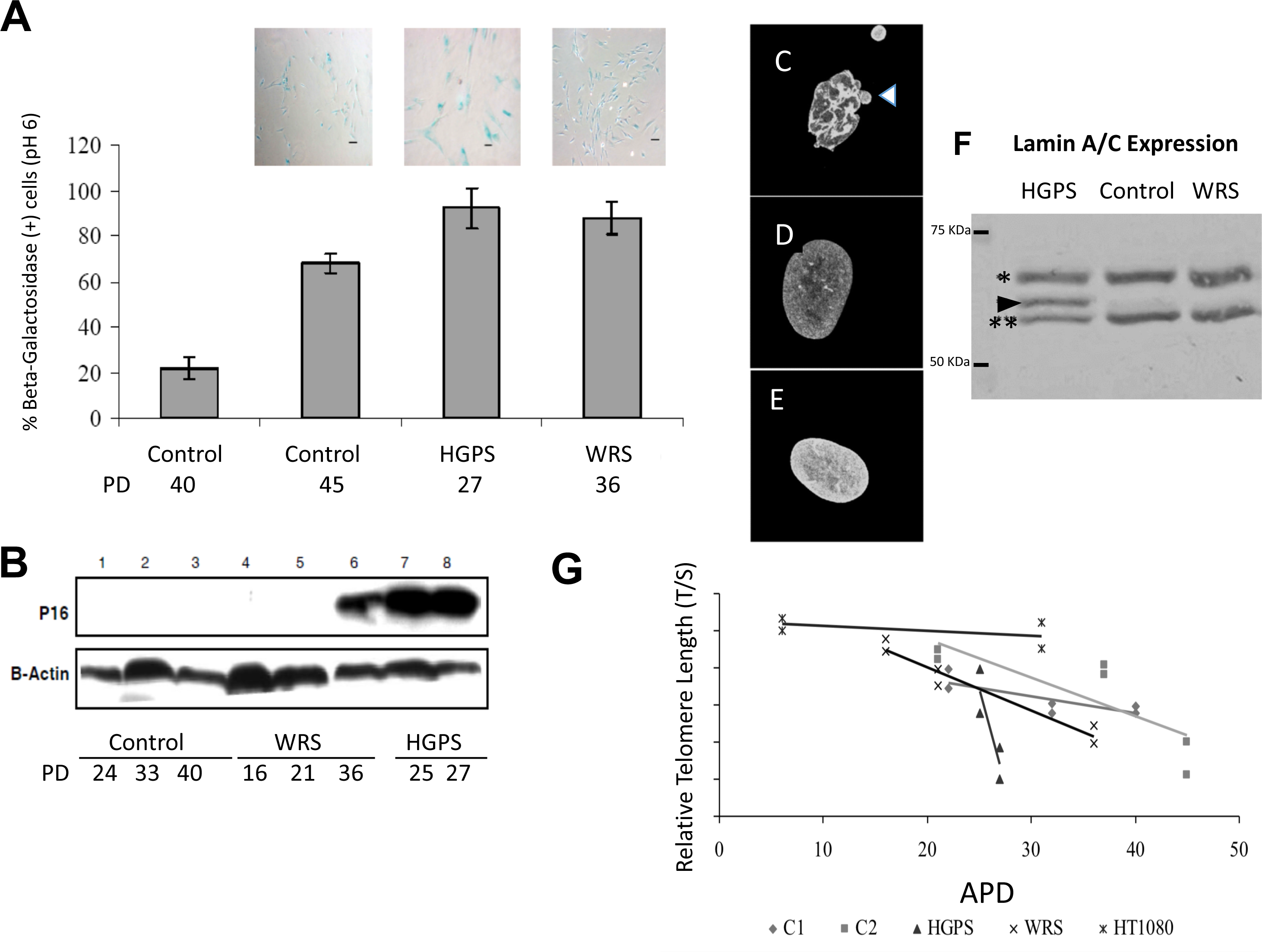
WRS fibroblasts are induced to early senescence. Senescence markers were analyzed at different population doublings (PD). (A) Beta-galactosidase staining (pH 6) was evident at PD 45 in controls, at PD 27 in HGPS and at PD 36 in WRS fibroblasts. (B) Expression of P16 was initially evident at PD 36 in WRS and at PD 25 in WRS. Expression of P16 was not found in controls cells up to PD 40. Nuclear morphology by using lamin A immunostaining is shown for (C) HGPS, (D) Control and (E) WRS fibroblasts cultures. Note that HGPS nuclei showed aberrant nuclear blebs (arrowhead). (F) Expression of Lamin A in HGP, WRS and control cells. WRS cells as well as control cells do not showed expression of progerin (arrowhead: 66 KDa) as observed in HGPS [*: Lamin A (70 KDa); ***: Lamin C (62 kDa)]. (G) Relative telomere length. The HT1080 osterosarcoma cell line (cross and circle) was used a positive control for telomere maintenance due to expression of telomerase. HGPS cells (triangles) show a sharp decrease in telomere length, while WRS cells (crosses) show a pattern of telomere shortening similar to control cells (squares and rhomboids) but starting at earlier population doublings (PD). APD: accumulated population doublings.

Nuclear morphology of HGPS (C), control (D) and WRS (E) cells was analyzed by Lamin A staining. HGPS nucleus shows the typical alterations in nuclear morphology, such as nuclear deformities and membrane blebbing (Figure 7C), as has been previously described. Nuclear morphology of WRS cells is very similar to control cells, with no apparent nuclear alterations (Figure 7D and E). To study the protein expression of different Lamin A and C isoforms, we used a polyclonal primary antibody that recognizes the region bordering aspartic acid 238 of lamin A, lamin C, and progerin. Expression of lamin A and C was observed in the different cell lines (Asterisks, Figure 7F). Progerin expression was present only in HGPS extracts (Figure 1F, arrowhead) as previously described (Scaffidi and Misteli, 2005), and no progerin expression was detected in control or WRS fibroblasts.

WRS cells showed a progressive decrease in the relative telomere length (Figure 7G, Blue line) with a rate comparable to control cells (Figure 7G, Orange line) but starting at earlier PD. In contrast, telomere length decrease in HGPS cells was very sharp and quickly (Figure 7G, Red line). The HT1080 osterosarcoma cell line (Figure 7G, Purple line) was used a positive control for telomere maintenance due to expression of telomerase.

## 4. DISCUSSION

Recently, bialelic heterozygous mutations in POLR3A have been described as the genetic cause of WRS, a rare syndrome with dramatic progeroid features present at birth (Jay et al., 2016; Paolacci et al., 2018). POLR3A encodes the main catalytic subunit of POLR3, which is important in the nucleolar homeostasis, ribosome biogenesis, protein translation and cell metabolism (Han et al., 2018; Tiku and Antebi, 2018; Turowski and Tollervey, 2016). WRS-derived fibroblasts analyzed in the present study carry a single pathogenic POLR3A variant c.3772_3773delCT (p.Leu1258Glyfs*12)]; and, although the second mutation is still unknown and, if present, most likely affects intronic regions or regulatory motifs of the gene (Paolacci et al., 2018), that clearly affects the expression level and compartmentalization of POLR3A in WRS fibroblasts.

In yeast, POLR3 biogenesis takes place in the cytoplasm where different POLR3 sub-units are assembled and then delivered into the nucleus by interaction with diverse chaperone proteins, where it performs its normal transcriptional activity (Han et al., 2018; Lesniewska and Boguta, 2017; Willis and Moir, 2018). The present mutation in POL3A caused a deletion of 5 nucleotides, leading to changes in the reading frame and generation of a premature stop codon and hence a truncated protein at the C-terminal region. The C-terminal POLR3A fragment lost in the present WRS case is important for interaction of POLR3A with additional POLR3 sub-units and also for its proper nuclear import. It has been described that mutations in POLR3A alter its capacity to interact with DNA and therefore cause a drastic alteration of its normal transcriptional function (Bernard et al., 2011; Paolacci et al., 2018), as is also observed in the present analysis. This suggests that the truncated POLR3A protein overexpress in the present case is in fact altering the normal transcriptional activity of POLR3.

Recently, increased function of POLR3A had gain interest due to its potential oncogenic function by increasing the levels of tRNA^Met^ and rRNAs (Arimbasseri and Maraia, 2016; Finlay-Schultz et al., 2017; Marshall and White, 2008). In addition, diverse genetic disorders are associated to mutations in POLR3A, in particular diseases with alterations in the development of cerebral white matter or leukodystrophies, including the 4H syndrome (hypomyelination, hypodontia and hypogonadotropic hypogonadism) (Bernard et al., 2011; Daoud et al., 2013; Terao et al., 2012). Most POLR3A mutations observed in WRS and leukodystrophy syndromes are compound heterozygous, and therefore the genotype-phenotype correlations must rely on the remaining level of functional POLR3A and on additional genetic or epigenetics changes not currently known. For instance, in a cellular model of POLR3A compound-heterozygous mutation related to hypomyelinating leukodystrophy, there was a decrease in POLR3A expression associated to reduced of brain cytoplasmic BC200 RNA (BCYRN1), which is involved in the regulation of translation of myelin basic protein (MBP), and therefore has a potential role in oligodendrocyte biology and related diseases (Choquet et al., 2019).

The cellular and molecular characteristics of WRS fibroblasts suggest that they undergo an early entrance into cell senescence, associated to a slightly larger nuclei and an increase in the number and area of nucleoli. WRS syndrome, which is considered a developmental disorder with aging characteristics, POLR3 function seems to be a key determinant of proper cellular development and its decrease/absence causes severe development alterations, including a premature senescence/aging phenotype with early lethality (Paolacci et al., 2017; Paolacci et al., 2018). In the present study, we demonstrate that fibroblast from a WRS patient have a sharp increase in the expression of a mutated version of POLR3A (∼142 KDa). The abnormal protein has atypical sub-cellular compartmentalization, having an increased in the intensity of the nuclear staining. This observations suggests that in WRS cells the normal function of POLR3A may be compromise in different ways: 1) abnormal subcellular localization due to alteration of its normal transport from the cytosol to the nucleus; and 2) increase in the levels and ectopic function of the mutated isoform present in WRS cells that may be translated into a dominant negative effect (Herskowitz, 1987), that behave as an apparent loss-of-function homozygote in a heterozygote (Veitia, 2007).

In HGPS, Lamin A mutations have been described to induce broad changes in nuclear architecture that impact on DNA dynamics and function, which are considered the basis for the early senescence and the phenotypic characteristics of this progeria (Broers et al., 2006; Burla et al., 2018; Burla et al., 2016; De Sandre-Giovannoli et al., 2003; Decker et al., 2009; Goldman et al., 2004; Mounkes and Stewart, 2004; Wheaton et al., 2017). Alterations in nuclear architecture have also been described in additional atypical progeroid individuals as well as in normal aging (Kudlow and Kennedy, 2006; McClintock et al., 2007; Scaffidi and Misteli, 2006). Although the cellular events that control nuclear size remain largely unknown, different evidence suggests that lamin proteins may be important contributors to this phenotype. For instance, studies in yeast suggest that a non-diffusible factor determinant for nuclear size relies on the relative quantity of cytoplasm around the nucleus, and possibly is lodged in the nucleus or in the RER (Neumann and Nurse, 2007). In Drosophila, proteins related to lamins (with no known human homologue) determine the size of the nucleus (Brandt et al., 2006). In Xenopus eggs the overexpression of prenylated forms of lamin B1, lamin B2 (Almonacid et al., 2018; Hutchison, 2002; Mukherjee et al., 2016) lead to nuclear membrane biogenesis, while interference with the normal-expression of lamins generates nuclei of reduced size (Hutchison, 2002). However, as Lamin A and C expression seem to be normal in WRS cells, the most plausible explanation for the nuclear phenotype observed maybe the broad alteration in cellular metabolism, in particular protein and ribosomal synthesis, associated to the alteration of POLR3 function.

Nucleoli are important for correct and rapid production and assembly of small and large ribosomal subunits into ribosomes, which is key for proper protein synthesis and adequate cell growth, proliferation and metabolism (Bahadori et al., 2018; Hernandez-Verdun et al., 2010), and its role in cell senescence and aging has become an important area of interest (Bahadori et al., 2018; Tiku and Antebi, 2018). The number and size of nucleoli varies depending on: 1) the metabolic demands of a particular cell such that cells with a high demand of protein synthesis, as pluripotent and cancer cells, have 1 to 2 big nucleoli (Boulon et al., 2010; Takada and Kurisaki, 2015); and 2) age of the cell, such that young yeast have 1 to 2 nucleoli, while older cells have 4 or more small nucleoli (Sinclair and Guarente, 1997). HGPS fibroblasts have a widespread increased protein turnover associated to higher ribosome biogenesis and larger nucleoli area as compared to control cells; and cells transfected to induce an increase expression of progerin have also larger nucleolar area and increased number of nucleoli (Buchwalter and Hetzer, 2017). These alterations in the nucleolus of HGPS must be cause directly by progerin-driven alterations in heterochromatin function, inducing an increased in transcription of rDNA promoters and thus causing increase in the expression of rRNA and number of nucleolus (Buchwalter and Hetzer, 2017). Interestingly, the increase in nucleolar area and rRNA observed in HGPS are directly correlated to donor age of primary fibroblasts (Buchwalter and Hetzer, 2017; Takada and Kurisaki, 2015; Tsekrekou et al., 2017). Primary WRS fibroblasts also have an increase in the number and total area of nucleoli, most probably related to a direct impact of POLR3 alteration upon nucleolar function. Supporting these observations, POLR3A-related Leukodystrophies showed abnormal ribosome regulation and reduction in protein synthesis, which are important hallmarks of nucleolar dysfunction (Dorboz et al., 2018; Tetreault et al., 2011). In a recent report, the expression of the CBX4 protein, part of the polycomb repressive complex, was found reduced during senescence of human Mesenchymal Stem Cells (hMSC); while overexpression of CBX4 rescued the senescence phenotype of hMSC in a manner dependent on maintenance of nucleolar function (Ren et al., 2019). In addition, decrease expression of fibrillarin in hMSC was also associated to decrease in cell proliferation, induction of cell senescence and increase in the nucleolar area (Ren et al., 2019).

Nucleolar stress such as nucleolar disruption/fragmentation or alterations in ribosomal biogenesis is widely accepted as a central regulator of P53 stabilization and activity, as proposed by the Rubbi and Milner model (Rubbi and Milner, 2003), which in in turn leads to P53-mediated cell senescence (Pinho et al., 2019). This is currently known as impaired ribosome biogenesis checkpoint (IRBC) (Gentilella et al., 2017), in which the nucleolus becomes an important stress sensor responsible for the activation of P53 (Boulon et al., 2010). The IRBC involves the translocation from the nucleolus to the nucleoplasm of diverse proteins such as p14ARF, ribosomal proteins (RP: RP5 and RP11) and 5S rRNA, that interact and inhibit MDM2 and Hdm2, leading to activation of P53 (Boulon et al., 2010; Gjerset and Bandyopadhyay, 2006; Turi et al., 2019). WRS fibroblasts have a clear increase in pP53 expression as compared to control fibroblasts, suggesting a potential pathway of POLR3-mediated P53 regulation that is lost upon POLR3A mutation. As mentioned, the regulation of P53 is mediated in part by 5S rRNA, which is a specific POLR3 target gen (Onofrillo et al., 2017). WRS cells have a high increase of H2AX activity. H2AX is involved in the recognition of DNA damage and the initiation of a DNA damage response (DDR). Recently, control of genomic stability has been linked to the normal function of the nucleolus, and disruption of the nucleolus is associated to increase in phosphorylation of H2AX (Keil et al., 2019). However, the exact mechanism of POLR3A-mediated P53 and H2AX activation remains to be analyzed in the WRS context.

POLR3 activity is positively controlled by the Target of Rapamycin Kinase Complex I (TORC1) a key driver of cellular anabolism and a longevity determinant, such that a reduction in POLR3 levels extends chronological lifespan in worms, flies and yeast (Filer et al., 2017). The apparent contradictory observations of POLR3 function related to longevity and to a progeroid syndrome such as WRS may be explained by the optimized role of POLR3 for growth and reproductive fitness early during development and being most likely detrimental for organismal lifespan (Filer et al., 2017), constituting a potential example of antagonistic pleiotropy, an important hypothesis of human aging (Austad and Hoffman, 2018).

## 5. CONCLUSION

The present WRS cells undergo a process of premature cell senescence, associated to diverse changes: increase expression of mutated POLR3A associated to alteration of its transcriptional function, increase in the number and area of the nucleolus, increased DNA damage and telomere shortening, activation of P53, changes that maybe causing a defect in ribosome biogenesis and protein translation (Figure 8). These observations open new avenues to analyze POLR3A involvement in potential pathways that might explain its role in cell senescence and human aging. Particularly interesting is the use of this WRS model to unravel key roles of the nucleolus in the physiology of aging. To better understand all these processes, it is important to perform a comparative analysis of the transcriptional profile in WRS and control cells. We believe that the morphological and molecular characteristics of WRS cells are important for proper diagnosis and classification of WRS, and further contribute to the genotype-phenotype correlations of POLR3A mutations.

**Figure 8.**
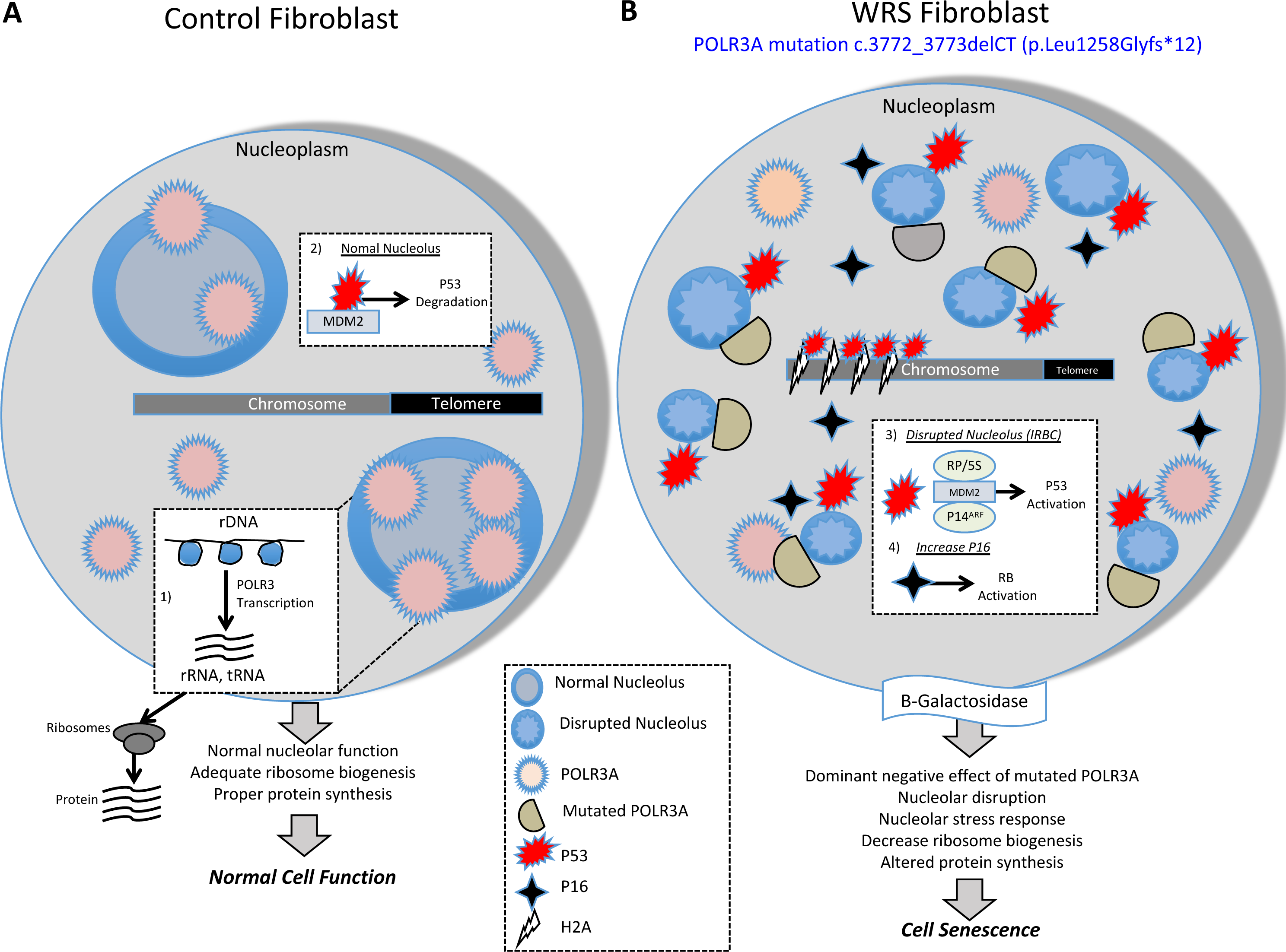
Model proposed for the effect of POLR3A mutations in WRS fibroblasts nucleoplasm. (A) In control fibroblasts POLR3A is mainly localized in the nucleoplasm, which is associated to normal number and function of nucleolus and normal telomere length. In a normal nucleolus: 1) rDNA is transcribed by POLR3 to produced rRNA, tRNA, and other RNAs, which are important for proper ribosome biogenesis and protein translation; 2) P53 binding to MDM2 induces its proteosomal degradation and keeps p53 inhibited. (B) WRS fibroblasts with a compound heterozygous POLR3A mutation c.3772_3773delCT (p.Leu1258Glyfs*12) have increase expression of mutant protein and reduction in expression of normal POLR3A in the nucleoplasm. This alteration in POLR3A is associated to disruption/stress and increased number of nucleoli; decrease telemore length, increased in DNA damage (increase pP53 and pH2AX expression), increased in expression of P16 and beta-glactosidase. In addition, disruption of the nucleolus: 3) released P14^ARF^ to the nucleoplasm where it binds and inhibits MDM2 leading to stabilization and activation of P53; 4) Increased P16 that leads to inhibition of CDKs-cyclin and activation of Rb protein. Overall, nucleolar dysfunction is associated to abnormal ribosomal biogenesis and protein synthesis that are associated to a senescence phenotype.

## ACKNOWLEDGEMENTS

This work was supported by grants from DIB-Universidad Nacional de Colombia and COLCIENCIAS. We gratefully acknowledge Dr. Gabriel Gutierrez-Ospina’s advice and support during N.R.’s short visit to his lab (Mobility program fellowship 2008, “Red de Macro Universidades de America Latina y el Caribe”).

## AUTHORS CONTRIBUTIONS

CT B and EV-R: carried out POLR3A studies, nuclear and nucleolus analysis and confocal microcopy analysis; NJR-S: carried out cell sencescence studies and nuclear size quantification; CG: carried out telomere lenght studies; LO-C and GG-O: carried out western blot analysis for Lamin A expression and instructed NJR-S; GGO: provided input and logistic for cell culture experiments, and provide continuous scientific feedback during NJR-S short-stay in his laboratory; AS-H: performed protein structural analysis; CA-B: conceived of the study, and participated in its design and helped to draft the manuscript; HA: conceived of the study, and participated in its design and coordination and helped to draft the manuscript; GA: conceived of the study, and participated in its design and coordination and wrote the manuscript with input from CA-B, HA, GGO, NJR-S.

## REFERENCES

Ahmed, M.S., Ikram, S., Bibi, N., Mir, A., 2018. Hutchinson-Gilford Progeria Syndrome: A Premature Aging Disease. Mol Neurobiol 55, 4417–4427.

Almonacid, M., Terret, M.E., Verlhac, M.H., 2018. Control of nucleus positioning in mouse oocytes. Semin Cell Dev Biol 82, 34–40.

Arboleda, G., Morales, L.C., Quintero, L., Arboleda, H., 2011. Neonatal progeroid syndrome (Wiedemann-Rautenstrauch syndrome): report of three affected sibs. Am J Med Genet A 155A, 1712–1715.

Arboleda, H., Arboleda, G., 2005. Follow-up study of Wiedemann-Rautenstrauch syndrome: long-term survival and comparison with Rautenstrauch’s patient “G”. Birth Defects Res A Clin Mol Teratol 73, 562–568.

Arimbasseri, A.G., Maraia, R.J., 2016. RNA Polymerase III Advances: Structural and tRNA Functional Views. Trends Biochem Sci 41, 546–559.

Austad, S.N., Hoffman, J.M., 2018. Is antagonistic pleiotropy ubiquitous in aging biology? Evol Med Public Health 2018, 287–294.

Bahadori, M., Azizi, M.H., Dabiri, S., 2018. Recent Advances on Nucleolar Functions in Health and Disease. Arch Iran Med 21, 600–607.

Baran, V., Vesela, J., Rehak, P., Koppel, J., Flechon, J.E., 1995. Localization of fibrillarin and nucleolin in nucleoli of mouse preimplantation embryos. Mol Reprod Dev 40, 305–310.

Bernard, G., Chouery, E., Putorti, M.L., Tetreault, M., Takanohashi, A., Carosso, G., Clement, I., Boespflug-Tanguy, O., Rodriguez, D., Delague, V., Abou Ghoch, J., Jalkh, N., Dorboz, I., Fribourg, S., Teichmann, M., Megarbane, A., Schiffmann, R., Vanderver, A., Brais, B., 2011. Mutations of POLR3A encoding a catalytic subunit of RNA polymerase Pol III cause a recessive hypomyelinating leukodystrophy. Am J Hum Genet 89, 415–423.

Binz, N., Shalaby, T., Rivera, P., Shin-ya, K., Grotzer, M.A., 2005. Telomerase inhibition, telomere shortening, cell growth suppression and induction of apoptosis by telomestatin in childhood neuroblastoma cells. Eur J Cancer 41, 2873–2881.

Boulon, S., Westman, B.J., Hutten, S., Boisvert, F.M., Lamond, A.I., 2010. The nucleolus under stress. Mol Cell 40, 216–227.

Brandt, A., Papagiannouli, F., Wagner, N., Wilsch-Brauninger, M., Braun, M., Furlong, E.E., Loserth, S., Wenzl, C., Pilot, F., Vogt, N., Lecuit, T., Krohne, G., Grosshans, J., 2006. Developmental control of nuclear size and shape by Kugelkern and Kurzkern. Curr Biol 16, 543–552.

Broers, J.L., Ramaekers, F.C., Bonne, G., Yaou, R.B., Hutchison, C.J., 2006. Nuclear lamins: laminopathies and their role in premature ageing. Physiol Rev 86, 967–1008.

Buchwalter, A., Hetzer, M.W., 2017. Nucleolar expansion and elevated protein translation in premature aging. Nat Commun 8, 328.

Burla, R., La Torre, M., Merigliano, C., Verni, F., Saggio, I., 2018. Genomic instability and DNA replication defects in progeroid syndromes. Nucleus 9, 368–379.

Burla, R., La Torre, M., Saggio, I., 2016. Mammalian telomeres and their partnership with lamins. Nucleus 7, 187–202.

Canella, D., Praz, V., Reina, J.H., Cousin, P., Hernandez, N., 2010. Defining the RNA polymerase III transcriptome: Genome-wide localization of the RNA polymerase III transcription machinery in human cells. Genome Res 20, 710–721.

Cao, H., Hegele, R.A., 2003. LMNA is mutated in Hutchinson-Gilford progeria (MIM 176670) but not in Wiedemann-Rautenstrauch progeroid syndrome (MIM 264090). J Hum Genet 48, 271–274.

Cawthon, R.M., 2002. Telomere measurement by quantitative PCR. Nucleic Acids Res 30, e47.

Childs, B.G., Bussian, T.J., Baker, D.J., 2019. Cellular Identification and Quantification of Senescence-Associated beta-Galactosidase Activity In Vivo. Methods Mol Biol 1896, 31–38.

Choquet, K., Forget, D., Meloche, E., Dicaire, M.J., Bernard, G., Vanderver, A., Schiffmann, R., Fabian, M.R., Teichmann, M., Coulombe, B., Brais, B., Kleinman, C.L., 2019. Leukodystrophy-associated POLR3A mutations down-regulate the RNA polymerase III transcript and important regulatory RNA BC200. J Biol Chem 294, 7445–7459.

Csoka, A.B., Cao, H., Sammak, P.J., Constantinescu, D., Schatten, G.P., Hegele, R.A., 2004. Novel lamin A/C gene (LMNA) mutations in atypical progeroid syndromes. J Med Genet 41, 304–308.

Daoud, H., Tetreault, M., Gibson, W., Guerrero, K., Cohen, A., Gburek-Augustat, J., Synofzik, M., Brais, B., Stevens, C.A., Sanchez-Carpintero, R., Goizet, C., Naidu, S., Vanderver, A., Bernard, G., 2013. Mutations in POLR3A and POLR3B are a major cause of hypomyelinating leukodystrophies with or without dental abnormalities and/or hypogonadotropic hypogonadism. J Med Genet 50, 194–197.

De Sandre-Giovannoli, A., Bernard, R., Cau, P., Navarro, C., Amiel, J., Boccaccio, I., Lyonnet, S., Stewart, C.L., Munnich, A., Le Merrer, M., Levy, N., 2003. Lamin a truncation in Hutchinson-Gilford progeria. Science 300, 2055.

Decker, M.L., Chavez, E., Vulto, I., Lansdorp, P.M., 2009. Telomere length in Hutchinson-Gilford progeria syndrome. Mech Ageing Dev 130, 377–383.

Dimri, G.P., Lee, X., Basile, G., Acosta, M., Scott, G., Roskelley, C., Medrano, E.E., Linskens, M., Rubelj, I., Pereira-Smith, O., et al., 1995. A biomarker that identifies senescent human cells in culture and in aging skin in vivo. Proc Natl Acad Sci U S A 92, 9363–9367.

Dorboz, I., Dumay-Odelot, H., Boussaid, K., Bouyacoub, Y., Barreau, P., Samaan, S., Jmel, H., Eymard-Pierre, E., Cances, C., Bar, C., Poulat, A.L., Rousselle, C., Renaldo, F., Elmaleh-Berges, M., Teichmann, M., Boespflug-Tanguy, O., 2018. Mutation in POLR3K causes hypomyelinating leukodystrophy and abnormal ribosomal RNA regulation. Neurol Genet 4, e289.

Eriksson, M., Brown, W.T., Gordon, L.B., Glynn, M.W., Singer, J., Scott, L., Erdos, M.R., Robbins, C.M., Moses, T.Y., Berglund, P., Dutra, A., Pak, E., Durkin, S., Csoka, A.B., Boehnke, M., Glover, T.W., Collins, F.S., 2003. Recurrent de novo point mutations in lamin A cause Hutchinson-Gilford progeria syndrome. Nature 423, 293–298.

Filer, D., Thompson, M.A., Takhaveev, V., Dobson, A.J., Kotronaki, I., Green, J.W.M., Heinemann, M., Tullet, J.M.A., Alic, N., 2017. RNA polymerase III limits longevity downstream of TORC1. Nature 552, 263–267.

Finlay-Schultz, J., Gillen, A.E., Brechbuhl, H.M., Ivie, J.J., Matthews, S.B., Jacobsen, B.M., Bentley, D.L., Kabos, P., Sartorius, C.A., 2017. Breast Cancer Suppression by Progesterone Receptors Is Mediated by Their Modulation of Estrogen Receptors and RNA Polymerase III. Cancer Res 77, 4934–4946.

Gentilella, A., Moron-Duran, F.D., Fuentes, P., Zweig-Rocha, G., Riano-Canalias, F., Pelletier, J., Ruiz, M., Turon, G., Castano, J., Tauler, A., Bueno, C., Menendez, P., Kozma, S.C., Thomas, G., 2017. Autogenous Control of 5’TOP mRNA Stability by 40S Ribosomes. Mol Cell 67, 55–70 e54.

Gil, M.E., Coetzer, T.L., 2004a. Real-time quantitative PCR of telomere length. Mol Biotechnol 27, 169–172.

Gil, M.E., Coetzer, T.L., 2004b. Real-time quantitative RT-PCR for human telomere elongation reverse transcriptase in chronic myeloid leukemia. Leuk Res 28, 969–972.

Gjerset, R.A., Bandyopadhyay, K., 2006. Regulation of p14ARF through subnuclear compartmentalization. Cell Cycle 5, 686–690.

Goldman, R.D., Shumaker, D.K., Erdos, M.R., Eriksson, M., Goldman, A.E., Gordon, L.B., Gruenbaum, Y., Khuon, S., Mendez, M., Varga, R., Collins, F.S., 2004. Accumulation of mutant lamin A causes progressive changes in nuclear architecture in Hutchinson-Gilford progeria syndrome. Proc Natl Acad Sci U S A 101, 8963–8968.

Green, D.R., Kroemer, G., 2009. Cytoplasmic functions of the tumour suppressor p53. Nature 458, 1127–1130.

Han, Y., Yan, C., Fishbain, S., Ivanov, I., He, Y., 2018. Structural visualization of RNA polymerase III transcription machineries. Cell Discov 4, 40.

Hernandez-Verdun, D., Roussel, P., Thiry, M., Sirri, V., Lafontaine, D.L., 2010. The nucleolus: structure/function relationship in RNA metabolism. Wiley Interdiscip Rev RNA 1, 415–431.

Herskowitz, I., 1987. Functional inactivation of genes by dominant negative mutations. Nature 329, 219–222.

Hoffmann, N.A., Jakobi, A.J., Moreno-Morcillo, M., Glatt, S., Kosinski, J., Hagen, W.J., Sachse, C., Muller, C.W., 2015. Molecular structures of unbound and transcribing RNA polymerase III. Nature 528, 231–236.

Huang, S., Kennedy, B.K., Oshima, J., 2005. LMNA mutations in progeroid syndromes. Novartis Found Symp 264, 197–202; discussion 202-197, 227-130.

Hutchison, C.J., 2002. Lamins: building blocks or regulators of gene expression? Nat Rev Mol Cell Biol 3, 848–858.

Jay, A.M., Conway, R.L., Thiffault, I., Saunders, C., Farrow, E., Adams, J., Toriello, H.V., 2016. Neonatal progeriod syndrome associated with biallelic truncating variants in POLR3A. Am J Med Genet A 170, 3343–3346.

Keil, M., Meyer, M.T., Dannheisig, D.P., Maerz, L.D., Philipp, M., Pfister, A.S., 2019. Loss of Peter Pan protein is associated with cell cycle defects and apoptotic events. Biochimica et biophysica acta. Molecular cell research 1866, 882–895.

Khatter, H., Vorlander, M.K., Muller, C.W., 2017. RNA polymerase I and III: similar yet unique. Curr Opin Struct Biol 47, 88–94.

Kopp, B., Khoury, L., Audebert, M., 2019. Validation of the gammaH2AX biomarker for genotoxicity assessment: a review. Archives of toxicology 93, 2103–2114.

Kruiswijk, F., Labuschagne, C.F., Vousden, K.H., 2015. p53 in survival, death and metabolic health: a lifeguard with a licence to kill. Nat Rev Mol Cell Biol 16, 393–405.

Kudlow, B.A., Kennedy, B.K., 2006. Aging: progeria and the lamin connection. Curr Biol 16, R652–654.

Kuo, C.H., Wells, W.W., 1978. beta-Galactosidase from rat mammary gland. Its purification, properties, and role in the biosynthesis of 6beta-O-D-galactopyranosyl myo-inositol. J Biol Chem 253, 3550–3556.

Lesniewska, E., Boguta, M., 2017. Novel layers of RNA polymerase III control affecting tRNA gene transcription in eukaryotes. Open Biol 7.

MacLaine, N.J., Hupp, T.R., 2011. How phosphorylation controls p53. Cell Cycle 10, 916–921.

Marshall, L., White, R.J., 2008. Non-coding RNA production by RNA polymerase III is implicated in cancer. Nat Rev Cancer 8, 911–914.

Martin, G.M., 1982. Syndromes of accelerated aging. Natl Cancer Inst Monogr 60, 241–247.

McClintock, D., Ratner, D., Lokuge, M., Owens, D.M., Gordon, L.B., Collins, F.S., Djabali, K., 2007. The mutant form of lamin A that causes Hutchinson-Gilford progeria is a biomarker of cellular aging in human skin. PLoS One 2, e1269.

McNicholas, S., Potterton, E., Wilson, K.S., Noble, M.E., 2011. Presenting your structures: the CCP4mg molecular-graphics software. Acta crystallographica. Section D, Biological crystallography 67, 386–394.

Morales, L.C., Arboleda, G., Rodriguez, Y., Forero, D.A., Ramirez, N., Yunis, J.J., Arboleda, H., 2009. Absence of Lamin A/C gene mutations in four Wiedemann-Rautenstrauch syndrome patients. Am J Med Genet A 149A, 2695–2699.

Mounkes, L.C., Stewart, C.L., 2004. Aging and nuclear organization: lamins and progeria. Curr Opin Cell Biol 16, 322–327.

Mukherjee, R.N., Chen, P., Levy, D.L., 2016. Recent advances in understanding nuclear size and shape. Nucleus 7, 167–186.

Nemeth, A., Grummt, I., 2018. Dynamic regulation of nucleolar architecture. Curr Opin Cell Biol 52, 105–111.

Neumann, F.R., Nurse, P., 2007. Nuclear size control in fission yeast. J Cell Biol 179, 593–600.

Onofrillo, C., Galbiati, A., Montanaro, L., Derenzini, M., 2017. The pre-existing population of 5S rRNA effects p53 stabilization during ribosome biogenesis inhibition. Oncotarget 8, 4257–4267.

Ou, H.L., Schumacher, B., 2018. DNA damage responses and p53 in the aging process. Blood 131, 488–495.

Paolacci, S., Bertola, D., Franco, J., Mohammed, S., Tartaglia, M., Wollnik, B., Hennekam, R.C., 2017. Wiedemann-Rautenstrauch syndrome: A phenotype analysis. Am J Med Genet A.

Paolacci, S., Li, Y., Agolini, E., Bellacchio, E., Arboleda-Bustos, C.E., Carrero, D., Bertola, D., Al-Gazali, L., Alders, M., Altmuller, J., Arboleda, G., Beleggia, F., Bruselles, A., Ciolfi, A., Gillessen-Kaesbach, G., Krieg, T., Mohammed, S., Muller, C., Novelli, A., Ortega, J., Sandoval, A., Velasco, G., Yigit, G., Arboleda, H., Lopez-Otin, C., Wollnik, B., Tartaglia, M., Hennekam, R.C., 2018. Specific combinations of biallelic POLR3A variants cause Wiedemann-Rautenstrauch syndrome. J Med Genet 55, 837–846.

Piekarowicz, K., Machowska, M., Dzianisava, V., Rzepecki, R., 2019. Hutchinson-Gilford Progeria Syndrome-Current Status and Prospects for Gene Therapy Treatment. Cells 8.

Pinho, M., Macedo, J., Logarinho, E., Pereira, P.S., 2019. NOL12 repression induces nucleolar stress-driven cellular senescence and is associated with normative aging. Mol Cell Biol.

Ren, X., Hu, B., Song, M., Ding, Z., Dang, Y., Liu, Z., Zhang, W., Ji, Q., Ren, R., Ding, J., Chan, P., Jiang, C., Ye, K., Qu, J., Tang, F., Liu, G.H., 2019. Maintenance of Nucleolar Homeostasis by CBX4 Alleviates Senescence and Osteoarthritis. Cell Rep 26, 3643–3656 e3647.

Rodier, F., Campisi, J., Bhaumik, D., 2007. Two faces of p53: aging and tumor suppression. Nucleic Acids Res 35, 7475–7484.

Rubbi, C.P., Milner, J., 2003. Disruption of the nucleolus mediates stabilization of p53 in response to DNA damage and other stresses. EMBO J 22, 6068–6077.

Saitsu, H., Osaka, H., Sasaki, M., Takanashi, J., Hamada, K., Yamashita, A., Shibayama, H., Shiina, M., Kondo, Y., Nishiyama, K., Tsurusaki, Y., Miyake, N., Doi, H., Ogata, K., Inoue, K., Matsumoto, N., 2011. Mutations in POLR3A and POLR3B encoding RNA Polymerase III subunits cause an autosomal-recessive hypomyelinating leukoencephalopathy. Am J Hum Genet 89, 644–651.

Scaffidi, P., Misteli, T., 2005. Reversal of the cellular phenotype in the premature aging disease Hutchinson-Gilford progeria syndrome. Nat Med 11, 440–445.

Scaffidi, P., Misteli, T., 2006. Lamin A-dependent nuclear defects in human aging. Science 312, 1059–1063.

Shieh, S.Y., Ikeda, M., Taya, Y., Prives, C., 1997. DNA damage-induced phosphorylation of p53 alleviates inhibition by MDM2. Cell 91, 325–334.

Sinclair, D.A., Guarente, L., 1997. Extrachromosomal rDNA circles--a cause of aging in yeast. Cell 91, 1033–1042.

Sinha, J.K., Ghosh, S., Raghunath, M., 2014. Progeria: a rare genetic premature ageing disorder. Indian J Med Res 139, 667–674.

Takada, H., Kurisaki, A., 2015. Emerging roles of nucleolar and ribosomal proteins in cancer, development, and aging. Cell Mol Life Sci 72, 4015–4025.

Terao, Y., Saitsu, H., Segawa, M., Kondo, Y., Sakamoto, K., Matsumoto, N., Tsuji, S., Nomura, Y., 2012. Diffuse central hypomyelination presenting as 4H syndrome caused by compound heterozygous mutations in POLR3A encoding the catalytic subunit of polymerase III. J Neurol Sci 320, 102–105.

Tetreault, M., Choquet, K., Orcesi, S., Tonduti, D., Balottin, U., Teichmann, M., Fribourg, S., Schiffmann, R., Brais, B., Vanderver, A., Bernard, G., 2011. Recessive mutations in POLR3B, encoding the second largest subunit of Pol III, cause a rare hypomyelinating leukodystrophy. Am J Hum Genet 89, 652–655.

Tiku, V., Antebi, A., 2018. Nucleolar Function in Lifespan Regulation. Trends Cell Biol 28, 662–672.

Tsekrekou, M., Stratigi, K., Chatzinikolaou, G., 2017. The Nucleolus: In Genome Maintenance and Repair. Int J Mol Sci 18.

Turi, Z., Lacey, M., Mistrik, M., Moudry, P., 2019. Impaired ribosome biogenesis: mechanisms and relevance to cancer and aging. Aging (Albany NY) 11, 2512–2540.

Turowski, T.W., Tollervey, D., 2016. Transcription by RNA polymerase III: insights into mechanism and regulation. Biochem Soc Trans 44, 1367–1375.

Ullrich, N.J., Gordon, L.B., 2015. Hutchinson-Gilford progeria syndrome. Handb Clin Neurol 132, 249–264.

Vasilishina, A., Kropotov, A., Spivak, I., Bernadotte, A., 2019. Relative Human Telomere Length Quantification by Real-Time PCR. Methods Mol Biol 1896, 39–44.

Veitia, R.A., 2007. Exploring the molecular etiology of dominant-negative mutations. The Plant cell 19, 3843–3851.

Villa, A., Navarro-Galve, B., Bueno, C., Franco, S., Blasco, M.A., Martinez-Serrano, A., 2004. Long-term molecular and cellular stability of human neural stem cell lines. Exp Cell Res 294, 559–570.

Wheaton, K., Campuzano, D., Ma, W., Sheinis, M., Ho, B., Brown, G.W., Benchimol, S., 2017. Progerin-Induced Replication Stress Facilitates Premature Senescence in Hutchinson-Gilford Progeria Syndrome. Mol Cell Biol 37.

Willis, I.M., Moir, R.D., 2018. Signaling to and from the RNA Polymerase III Transcription and Processing Machinery. Annu Rev Biochem 87, 75–100.

